# The triadic structure of personality space in the house mouse resembles that of species with known personality badges; suggesting reciprocal hierarchies are an evolutionary motif with mixed strategies being more common

**DOI:** 10.1101/2022.03.13.484125

**Authors:** Oren Forkosh, Barry Sinervo

**Affiliations:** Department of Cognition and Brain Sciences, The Hebrew University of Jerusalem, Jerusalem, Israel; Department of Animal Sciences, The Hebrew University of Jerusalem, Rehovot, Israel; Department of Ecology and Evolutionary Biology, Coastal Biology Building, 130 McAllister Way, University of California, Santa Cruz, California, 95064, USA

**Author notes:** Unfortunately, Barry passed away from cancer on March 15^th^, 2021, before we could publish this manuscript. It has been a great honor and a privilege to work with him, and he will be greatly missed.

## Abstract

Differences in animal personalities are generally hard to measure - even in an animal as common as the house mouse. Yet, in some rare cases, nature has provided us with clear visual cues as to the nature of a specific individual. These so-called badges are the case for the side botched lizard, often referred to as the ‘rock-paper-scissor’ lizard. Here, we show that mice have behavioral archetypes that are similar to the triadic social dynamics of the lizards. We find analogs in territoriality, aggressiveness, and pro-social behavior linking ultra-dominant, dominant, and subordinate mice to the lizard’s orange, blue, and yellow morphs accordingly. Yet, unlike the lizards, most mice in the study displayed mixed strategies rather than fall into clear behavioral archetypes. In such a case, it makes less sense to have distinct visual cues to distinguish between personalities. The result implies that, although visual distinctions are infrequent in nature, having multiple behavioral archetypes might serve as an evolutionary motif. In addition, we find that mammals, like mice, might have circular hierarchies, similar to the lizards, where no single behavioral strategy outweighs others.

## Main text

Individual differences are an essential property of all living things. One important source of diversity is behavior - in many cases, two individuals will vary in response to similar events. For example, while one person might startle in response to sudden noise, another might remain mostly indifferent. Consistent behavioral differences in humans are often referred to as personality traits and have been studied extensively in psychology (1). The common model for personality is the big-five personality traits, which, as the name suggests, breaks a person’s personality into five continuous factors (2). These five factors are usually determined using a self-report questionnaire.

The study of personality in animals has also gained considerable popularity in recent years, partly due to improvements in our ability to track large groups of animals for prolonged periods (3–6). Yet for animals, the definition of personality is still disputed, which is reflected by the multitude of definitions and names it is referred to, such as temperament, behavioral syndromes, coping styles, or just predisposition (7). Part of the dispute around animal personality results from the difficulty in measuring it since, unlike humans, self-report questionnaires are, of course, out of the question.

### Mouse Personality Space

Recently, it has been shown how animal personalities can be inferred directly and objectively from high-dimensional natural behavioral space (8). Personality traits have two essential properties: they differ among individuals, while being stable over time and across contexts (7). Therefore, the mathematical formulation of a trait informed directly by these properties would capture the maximum behavioral variability among individuals while maintaining minimum variability within individuals over time (Figure 1A). While this method is not species-specific, it has been demonstrated on mice, one of the most common model animals (Figure 1B). Traits found this way (which are referred to as identity domains or IDs) were shown to be stable across social context, do not change with age, explain the variability in performance in classical tests, and significantly correlated with gene expression in various brain regions.

**Figure 1.**
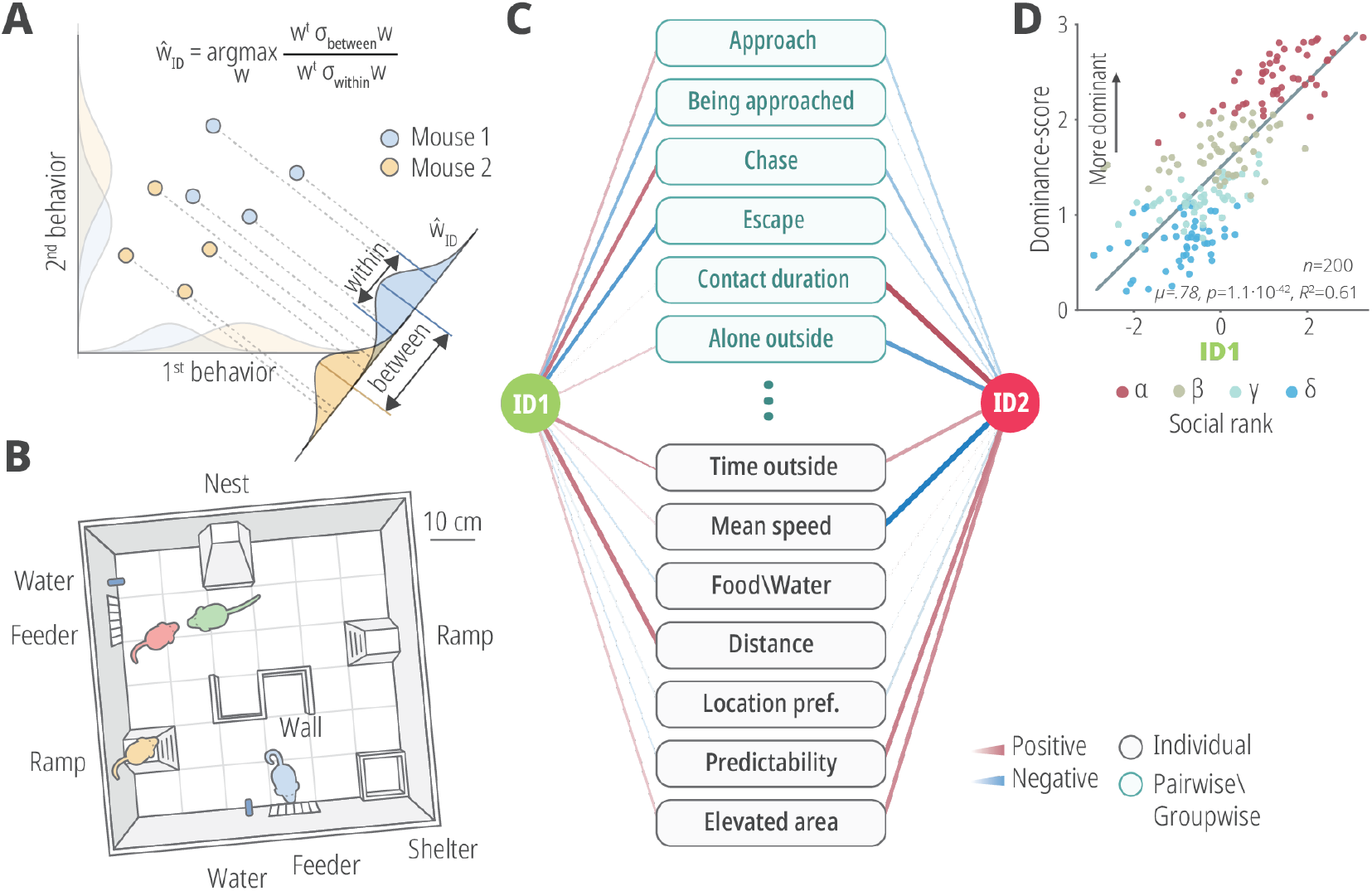
Inferring personality from behavior. (a) Personality traits both differ between individuals and are stable across time and context within the same individual. This definition of personality can be formulated as an optimization problem that is similar to linear discriminant analysis (LDA). (b) We measured the personalities of male mice housed in groups of four within an enriched arena. The mice spent four days within an arena which contained a closed nest, two feeders and two water bottles, two ramps, an S-shaped wall, and an open shelter. Each mouse was marked using a distinct hair dye, while his position and behavior were automatically tracked. (c) A total of four distinct personality traits were found, which were termed identity domains. The correlation between a subset of behaviors and the two most significant traits, ID1 and ID2 is shown here. (d) ID1 was significantly correlated with the dominance rank of each mouse, quantified here using the normalized David’s score.

A total of four stable traits, or identity domains (IDs), were found and labeled ID1 to ID4. The traits were ordered according to significance (using the Fisher-Rao criterion), and here we focus on the most significant traits, ID1 and ID2 (Figure 1C). The four identity domains were intentionally left unlabeled to avoid anthropomorphism. Yet, as each trait correlates with a distinct set of behaviors, we can use these behaviors to understand the traits. For example, ID1, the most significant trait, was found to correlate very significantly with the dominance hierarchy rank of each mouse (Figure 1D).

Yet, determining the identity domains of animals is not always feasible and often not even possible. In order to define the identity domains for mice, the trajectories of 200 mice within 50 groups were tracked continuously for at least four days within a controlled environment. Sixty unique behavioral readouts were automatically measured for each mouse, such as chases, approaches, and exploration. (9).

### Personality badges

However, on some relatively rare occasions, nature has “helped” by providing us clues to an animal’s expected behavior and personality. For example, in the cichlid fish Astatotilapia burtoni, non-territorial males are cryptically colored, in contrast to territorial males, which are either light blue or yellow (10). Moreover, the yellow males are more likely to become dominant when paired with blue conspecifics, although the coloring may change depending on their performance in aggressive interactions. Such so-called modifiable badges have been observed in other animals, including insects (11), birds (12, 13), lizards (14), and others.

In some cases, morphological cues remain persistent throughout the animal’s life span. For instance, white-throated sparrows have two morphs: white and tan, where the white males are usually more aggressive and provide less maternal care (15). Zebrafish selectively bred to be bold had elongated bodies and larger caudal regions compared to shy individuals (16).

One of the well-known examples of lifelong morphological differences is the side-blotched lizard (*Uta Stansburiana*), which is also often referred to as the rock-paper-scissors lizard (17). The males in this species have a triadic polymorphic variation in the color of their throat that also corresponds to distinct behavioral phenotypes (Figure 2A and Table 1). The orange-throated males are dubbed ultra-dominant since they tend to be bigger, more aggressive, maintain the largest territories, and have multiple females. Blue males have smaller territories and one or two females that they mate guard, but blue males have green-beard recognition and prefer to settle next to other blue males and cooperate against orange and yellow (18, 19). Against orange, one blue male will defend its (unrelated) partner and thus is altruistic in giving up its own fitness while preserving the recipient blue’s fitness. Against yellow, both blues benefit from cooperation and the relationship is thus mutualistic. Yellow males usually do not have a fixed territory or a fixed mate. Instead, the yellow males are attracted to the females of the orange lizards and mate with them while the orange males are away (taking advantage of the orange’s large territory). The yellow males mimic female behavior when confronted by territorial males. This genic behavior of blue also appears to structure a parallel rock-paper-scissors mating system in the European common lizard, *Zootoca vivipara* (20). These lizards are orange, white, and yellow, and the fitness of white males appears to be very high when other white males are common, indicative of evolutionary cooperation akin to the green beard of the side-blotched lizard, *Uta stansburiana*.

**Figure 2.**
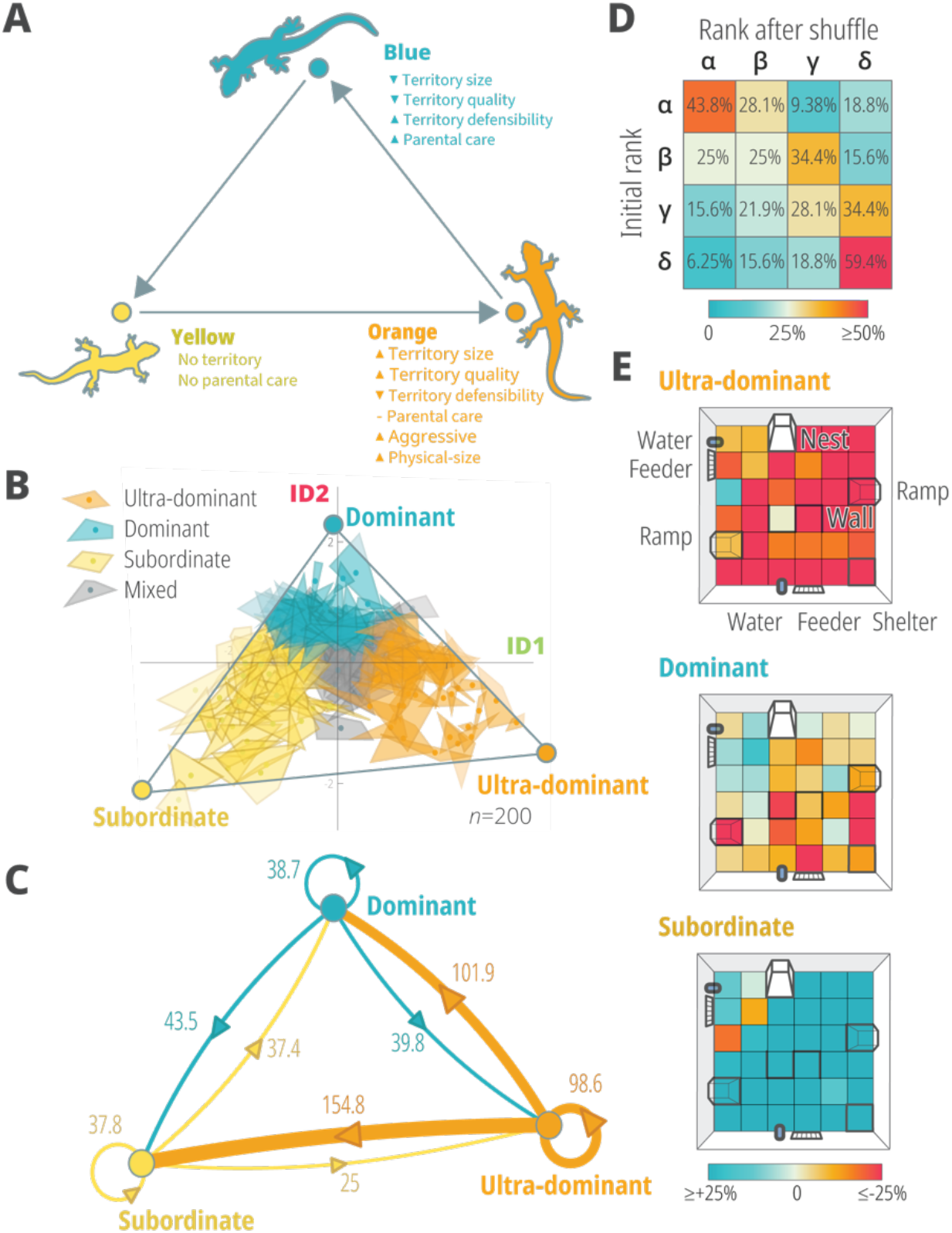
Three personality archetypes. (a) The males of the side-blotched lizard (Uta Stansburiana) have three known polymorphic variations in their throat color that coincide with distinct behavioral phenotypes. The dynamics between the three morphs, the orange, blue, and yellow variants, is often compared to the rock-paper-scissor game since the more aggressive orange males beat the less intrusive blue males, the territorial blue individuals usually fend off the territory-less yellows, while the opportunistic yellows mate with the females monopolized by the orange males. (b) The mice’s personality space spanned by the two most significant identity domains, ID1 and ID2, show three distinct behavioral strategies. (c) An average number of aggressive dyadic interactions, per day, between the three mouse archetypes. (d) Change in social rank following mice introduction into new social groups. (e) Territories of the mice according to the archetypes. The color represents the areas that the mice preferred or avoided compared to the average mouse.

**Table 1.**
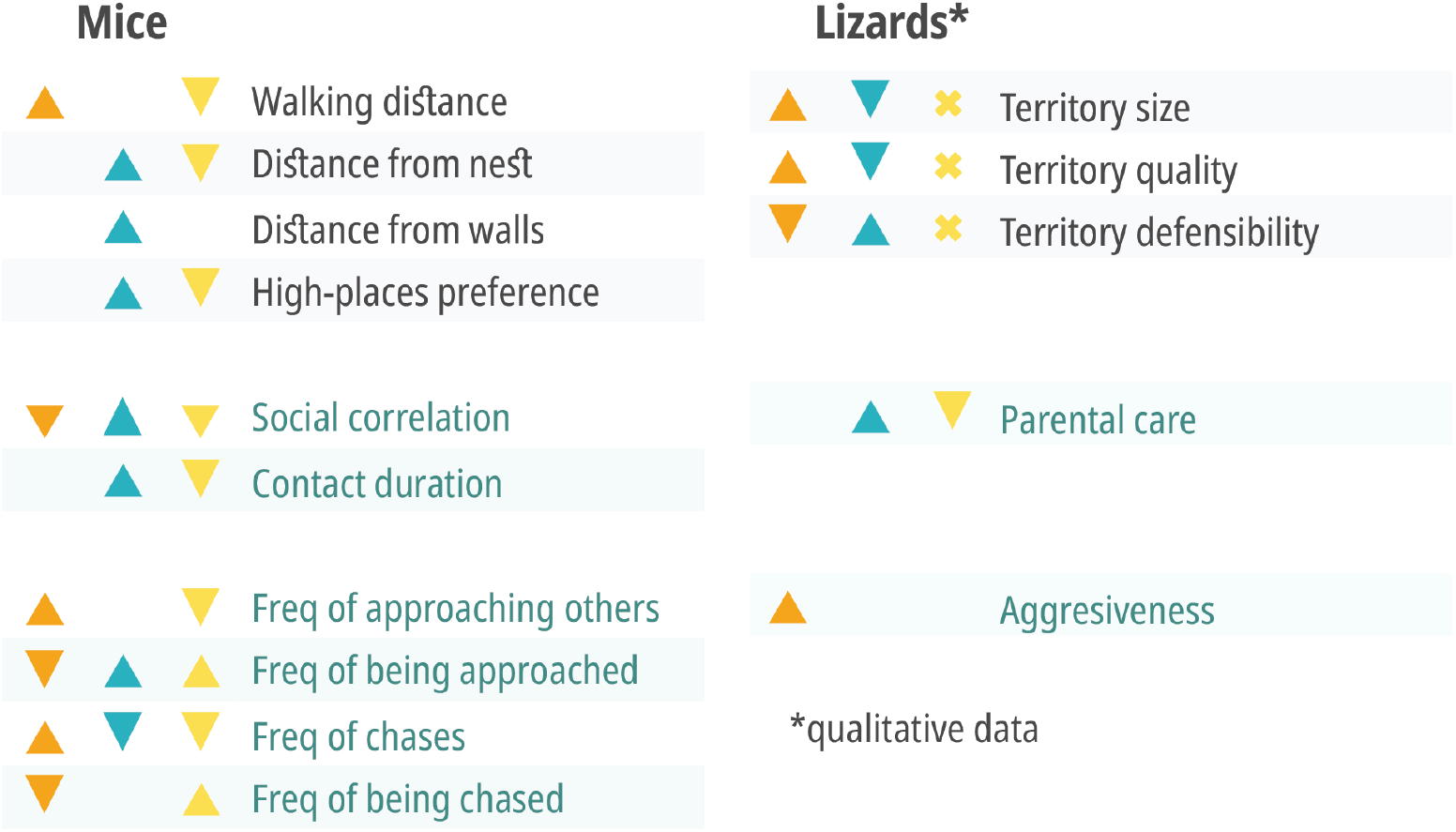
Behavioral differences between the archetypes. On the left, for the mice, an upward arrow represents a significance increase in comparison to the other two archetypes, while a downward arrow depicts significant decrease (with a Bonferroni correction to account for 60 behavioral readouts). The results for the lizards are qualitative results from various studies. The different colors correlate with the different archetypes: orange, blue, and yellow for the lizards, and accordingly ultra-dominant, dominant and subordinate for the mice. An ‘X’ in the lizard column marks the nonterritorial yellow morphs.

The nickname rock-paper-scissors given to these lizards comes from the interaction dynamics between the three morphs. Since orange lizards tend to be more aggressive, they will usually attack neighboring blue males in an agonistic interaction as the blue has a lower stamina (21) and eventually is ousted from its territory or dies, while its partner has time to mate and successfully produce progeny to the next generation. Blue females also prefer to settle with blue males (19). Similarly, blue males will usually beat yellow males because they assiduously defend their territory and their female(s) in green-beard mutualism. Yet, from an evolutionary ‘passing-your-genes’ perspective, because yellow males are able to reproduce with females monopolized by orange lizards, they are considered as having the upper hand in orange-yellow interactions. The number of offspring sired by each male morph of the lizards is intransitive and varies across social neighborhoods: when orange is common, yellow has the highest fitness, when yellow is common blue has most increased fitness and when blue is common, orange has the highest fitness (19, 22, 23).

## Methods

### Social boxes

Non-sibling male CD-1 (ICR) mice were housed in groups of four shortly after weaning (∼4 weeks). At around 7-8 weeks old, each mouse within a group was dyed using distinct hair colors. At the ages of 8-12 the mice were placed for at least four days within the enriched social boxes (35). The social boxes measured 60×60cms and included a large closed nest, an S-shaped wall, a small open shelter, as well as two feeders, two water bottles, and two elevated ramps (8). The mice were recorded using a lowlight overhead video camera which tracked them during the 12h dark cycle at a luminance level of 2 lux (full moon). From the video recordings, we extracted the position of each mouse, as well as 60 behavioral readouts such as chases (9), exploration, time in the feeder, synchronicity, etc. These behaviors were used to infer the identity domains of each mouse (8). A total of 200 mice within 50 groups were tracked in this way. See supplementary information for more details on the tracking system and behavioral analysis.

### Personality archetypes

To uncover the structure of personality space, we fitted a polytope to engulf the identity domains of all 200 mice (36). Using an unmixing algorithm, we found that a 3-vertice polytope was a significant fit to the personality space spanned by ID1 and ID2 (8). Here, we used the minimal volume simplex analysis, which is suitable for relatively small datasets since it does not allow for outliers. The vertices of the polytype correspond to extreme personality traits.

We divided the mice into three equally sized groups based on their nearest archetype (using the Euclidean metric). Each group included 49 mice, as we chose the largest possible group size such that each mouse was assigned a single archetype while keeping all groups the same size. Mice not assigned a specific archetype were labeled as having a ‘mixed strategy’ (n=53).

## Results

Interestingly, when looking at mice’s personality space, we can also identify three distinct behavioral strategies (Figure 2B). When looking at the significant two IDs, we obtained a personality space that is triangularly shaped. Inspired from the idea of Pareto optimality (24), the unique structure of personality space allows us to recognize three extreme behavioral archetypes which are captured by the vertices of the triangle. Since ID1 is related to hierarchy, we labeled the archetypes accordingly: ultra-dominant, dominant, and subordinate. We identified each mouse according to the closest archetype, while mice that overlapped with more than one archetype were referred to as mixed (see methods).

Somewhat surprising is that the aggressiveness levels of dominant mice towards the subordinates were not significantly different from the other way round (Figure 2C). But subordinate mice get their lower dominance scores because of the increased aggressiveness (in absolute numbers) directed from the ultra-dominant mice. This increase in aggressiveness does not depend on the group or the personality of the alpha male, since we find that subordinate mice tend to maintain their low rank even when placed in newly randomly formed groups (Figure 2D). The dominances’ behavior also tends to be more pro-social (Table 1). They showed higher levels of synchronicity, which is reflected by being positively correlated with others in the times they spend inside or outside the nest. Their social interactions are also usually longer and with fewer aggressive outcomes.

In terms of territoriality, dominant mice spend more time at the part of the arena which is furthest away from the safety of the nest. This area is, at the most part, occluded from the nest due to the wall and ramps (Figure 2E). In contrast, subordinate mice rarely exit the nest and usually remain in its close proximity limiting their outings for acquiring food and water. While the dominant and subordinate mice have distinct and disjoint locations, ultra-dominant mice, in general, spend more time outside of the nest and usually cover the entire area of the arena.

## Discussion

In several animal species, including lizards, fish, and others, visual differences might help determine social roles and thus decrease aggressiveness and prevent wasteful competition. But behavioral differences do not often manifest themselves with such distinct morphological variations. We know from human studies that personality traits can be subtly mediated through features like eye size (25) or facial symmetry (26). But these cues might be due to similarities to facial expressions we associate with certain personalities. It might be that in cases where the personalities themselves are not distinct, advertising your personality to others is less beneficial. And in some cases, it might actually be disadvantageous to let others know your true character, such as in cheating strategies. Another possibility is that mice use other modalities to advertise their personality, such as olfaction.

In a recent study of animal personalities, we found mice to have a personality space bounded by three distinct archetypes (8). While the mice personalities spanned the entire area of the triangle resulting in many ‘mixed strategies’, the archetypes we found bare several similarities to the well-known rock-paper-scissor lizards.

The dominant mice, like the blue lizards, had smaller territories, and of lesser quality (due to their distance from the nest), but are more defendable due to the presence of the elevated ramps and the s-shaped wall. The mice also tended to be more pro-social with higher social synchronicity and contact durations, which might be of more similarity to the monogamous nature of the blue lizards. The pro-sociality is akin to a rock-paper-scissors model of rodent mating systems in which promiscuous species occur in 3 types of territoriality: polygynous males with multiple males, monogamous males with a single female, but the monogamous males are more likely to care for progeny and sneaker males that lack territories and float about the matrix of polygynous and monogamous types (27). Thus, monogamous rodent males have genic care, akin to the genic (green-bearded) altruism of blue lizards. The ultra-dominant individuals, on the other hand (much like the orange lizards), were more aggressive and had a territory that spend the entire area of the arena. The subordinate mice, while as aggressive as the dominants, spent more time under the safety of the nest and kept exists short and dedicated mostly for accessing recourses such as food and water. They were also the main target for attacks by the ultra-dominant.

While we named the three archetypes according to their dominance rank, given the distinct behavioral differences it might be more accurate to relabel the ultra-dominant, dominant, and subordinate mice as aggressive, communal, and shy\timid accordingly. More work is needed to determine if there is indeed a rock-paper-scissor-like dynamic between mice archetypes. We need to examine the behavior of mice in a more natural setting while also including females in the study. Yet one of the main claims for the benefit of dominance is reproductive success. And while we do find some evidence linking dominance with the probability of siring offspring, the relation between the two is usually not very strong (28–30). This might imply that circular hierarchies, where no single behavioral strategy outweighs others, are more common in nature than previously thought.

For example, additional studies on pro-social behavior like paternal care by monogamous — dominant males — require analysis, as does analysis of dominant male personality types to neighboring dominant male personality types. The genic model of RPS for rodent mating systems ascribes dear enemy recognition between monogamous males that may be genic recognition of their own progeny extended to other dominant males with which they share the green-beard alleles for signal recognition and cooperation. A likely candidate for this green-beard is the Vasopressin receptor, V1aR which evolves mutations when rodents evolve from the promiscuous types of males to purely polygynous types of males. Sinervo et al. (27) also found that purely monogamous lineages gave rise to polygynous lineages at a high rate. These evolutionary transitions between monogamy and polygyny in rodents are linked to a point mutation in a Vasopressin receptor gene [V1aR: (31–33)]. V1aR is also linked to increased paternal behavior and reduced aggression (i.e., affiliative behavior) in monogamous species (34). The behaviors in the current study are consistent with this theory or rock-paper-scissors rodent mating systems.

